# Domain-Specific EPR Spectroscopy to Monitor Facilitated Dissociation of a DNA-Transcription Factor Complex

**DOI:** 10.1101/2025.06.13.659481

**Authors:** Ludovica Martina Epasto, Mahdi Khalil, Giuseppe Sicoli, Dennis Kurzbach

**Affiliations:** Faculty of Chemistry, Institute for Biological Chemistry, University of Vienna, Währinger Straße 38, 1090, Vienna, Austria; CNRS UMR 8516, LASIRE, University of Lille, C4 Building, Avenue Paul Langevin, 59655 Villeneuve d’Ascq Cedex, France

## Abstract

Transcription factors (TFs) regulate gene expression by interacting with specific DNA sequences in various manners, yet the mechanisms by which they dissociate from high-affinity DNA sites under physiological conditions remain not completely understood. Here, we employ continuous wave electron paramagnetic resonance (CW-EPR) spectroscopy in native solution state to resolve, with domain-level precision, the dissociation dynamics of the MYC-associated factor X (MAX) from its cognate EBOX DNA motif. Site-directed spin labeling reveals that in the absence of DNA, MAX undergoes a stepwise dissociation process—beginning with melting of the N-terminal disordered region (NTD), followed by the helix-loop-helix (HLH) domain, and culminating in leucine zipper (LZ) dimer dissociation. DNA binding reorganizes this process into a cooperative all-or-none transition, wherein destabilization of the LZ triggers concerted collapse of the entire trimeric MAX:MAX–DNA complex. Strikingly, the addition of a disordered BRCA1 fragment (residues 219–504) disrupts this cooperativity by selectively destabilizing DNA contacts in the NTD and HLH regions, without perturbing the LZ. This results in facilitated dissociation *via* competitive DNA binding, observed here directly at domain-level resolution. Our findings establish EPR spectroscopy as a uniquely sensitive tool for dissecting TF–DNA dynamics under physiological conditions, offering mechanistic insight into regulated unbinding processes inaccessible to ensemble methods.

## Introduction

Transcription factors (TFs) play central roles in regulating gene expression by targeting specific DNA motifs within chromatin^1–4^. MAX, the MYC-associated factor X, is a bHLH-LZ transcription factor that forms a homodimer in solution and recognizes the canonical enhancer box (EBOX) DNA element (CACGTG) in gene promoters.^5–9^ Max is a key player in regulating the transcriptional machinery, being involved in the control of a plethora of MYC-regulated genes. Indeed, disruption of MAX function has been linked to numerous oncogenic processes.^6, 10, 11^ While the thermodynamics and structural biology of MAX:MAX and MAX:MAX-DNA binding have been characterized, the kinetic and mechanistic basis of MAX binding to and dissociating from its cognate DNA remains poorly understood. In particular, it remains an open question of how TFs can execute their functions despite typically having nanomolar affinities to their cognate DNAs.^12, 13^

Facilitated dissociation—a mechanism wherein competitor proteins or nucleic acids actively promote TF release—has emerged as a plausible model for such regulation.^14–17^ Recently, it was shown that MAX undergoes large-scale dynamic fluctuations,^15^ which might underlie facilitated dissociation from or diffusion along DNA strands. However, direct domain-resolved experimental observations and descriptions of such events under near-physiological conditions have not yet been reported. Addressing this gap requires methodologies capable of resolving structural transitions in specific regions of a protein immediately as it dissociates from a DNA strand.

Here, we propose temperature-dependent continuous wave electron paramagnetic resonance (CW EPR) spectroscopy^18–25^ as a technique to track local conformational changes within individual domains of MAX upon DNA dissociation. By site-directed spin labeling at selected positions^26–28^ in the N-terminal DNA-binding domain, the helix-loop-helix motif, and the C-terminal leucine zipper, we map the domain-specific temperature-induced unfolding pathway of MAX *via* the stability of its individual components. We find that (i) the leucine zipper acts as a main anchor between both MAX strands and as a key stabilizing moiety, (ii) binding of cognate DNA to the N-terminal MAX domain stabilizes the homodimer leading to a cooperative DNA-MAX:MAX dissociation process, wherein the DNA-MAX and MAX-MAX interaction are simultaneously disrupted, (iii) the presence of the disordered BRCA1 fragment, housing its DNA-binding domain, promotes facilitated dissociation by destabilizing the DNA-bound form of MAX without perturbing the dimeric scaffold.

The sensitivity of EPR to changes in local spin label dynamics, combined with its applicability at near-physiological temperatures and concentrations^29, 30^, thus offered us a unique window into protein-DNA interactions that are difficult to access by conventional methods. Together with complementary NMR data, this work provides a mechanistic framework for transcription factor unbinding and highlights the utility of EPR for detecting complex biomolecular dissociation events with site specificity.

## Results

The bHLH-LZ transcription factor MAX forms a characteristic homodimer, in which each monomer comprises an N-terminal intrinsically disordered domain (NTD) that binds to DNA double-strands, a helix-loop-helix (HLH) motif, and a C-terminal coiled-coil leucine zipper (LZ) responsible for dimerization^7–9, 31^ (*Fig. 1., A*.). The NTD folds upon binding to DNA into another pair of helices embracing the nucleic acid ligand.^32^ The homodimer exists in a dynamic equilibrium with its monomeric forms whereas low concentrations and high temperatures promote dissociation of the dimer^26, 27, 33^, giving rise to a heterogeneous conformational ensemble.

**Figure 1.**
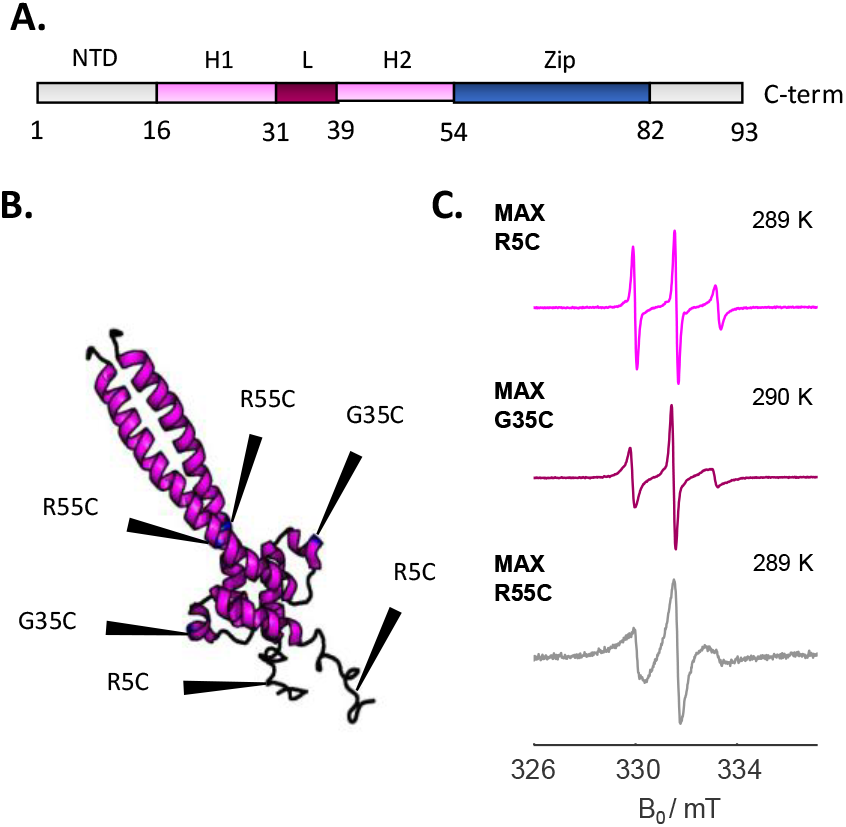
Spin labelled MAX in CW EPR. A. Schematic representation of MAX monomer structure. The two helices of the HLH domain are indicated in pink, while the leucine zipper domain is represented in blue. The N-and C-terminus are indicated in gray. B. MAX dimer structure in solution. The black arrows indicate the position of the residues mutated to cysteines for the spin labelling procedure. C. EPR spectra of MAX R5C, G35C, and R55C at temperatures below 300 K. The broadness of the spectra suggests the slow tumbling of the protein at low temperature; in this case, the spectra can be simulated by a single-component system.

To probe the dissociation of the MAX:MAX-DNA complex, we conducted temperature-dependent CW EPR spectroscopy on MAX variants site-specifically labeled with MTSL at three mutated residues: R5C (NTD), G35C (loop of HLH), and R55C (LZ) (*Fig. 1., B*.). These mutations do not perturb dimerization or ligand binding.^15, 28^

In the following, we will first describe our investigation of the neat MAX:MAX dimer to obtain reference data before moving on to the more complex MAX:MAX-DNA trimer. Finally, we will highlight how a competing BRCA1 ligand facilitates the dissociation process.

### Sequential unfolding of MAX

To probe the stability of the different motifs of the MAX:MAX dimer, we recorded temperature-dependent CW EPR data. The rationale behind this approach was to employ the dependence of MTSL CW EPR spectra on the rotational freedom of the spin label. Upon undergoing the well-documented temperature-induced transition of MAX from the rigid dimer to the dynamic intrinsically disordered state^13^, the MTSL labels were, thus, expected to lead to an abrupt change in EPR line shape.

At low temperatures (< 300 K), all EPR spectra of all three variants in the absence of DNA were well-described by a dominating single spectral component representing fast rotational motion with a rotational correlation time τ_c_ of 0.89 ns, 1.82 ns, and 3.98 ns for R5C, G35C, and R55C, respectively. Additionally, small (<10%) contributions of an even faster species were found. These spectra are representative of the MAX:MAX dimer dominating the conformational space at low temperatures (*Fig. 1, C*.).

Upon heating >300 K, a second, faster component suddenly emerged (as expected), increasing in population with temperature until it dominated the spectral signature (*Fig. 2, A* and *B*.). This biphasic spectral evolution reflects a redistribution within the conformational ensemble toward states with greater local mobility. Spectral simulations confirmed that a two-component model with two distinct rotational correlation times provided a minimal and robust fit to the data. In contrast, single-component models failed to reproduce the observed features (*Fig. 2, C*.).

**Figure 2.**
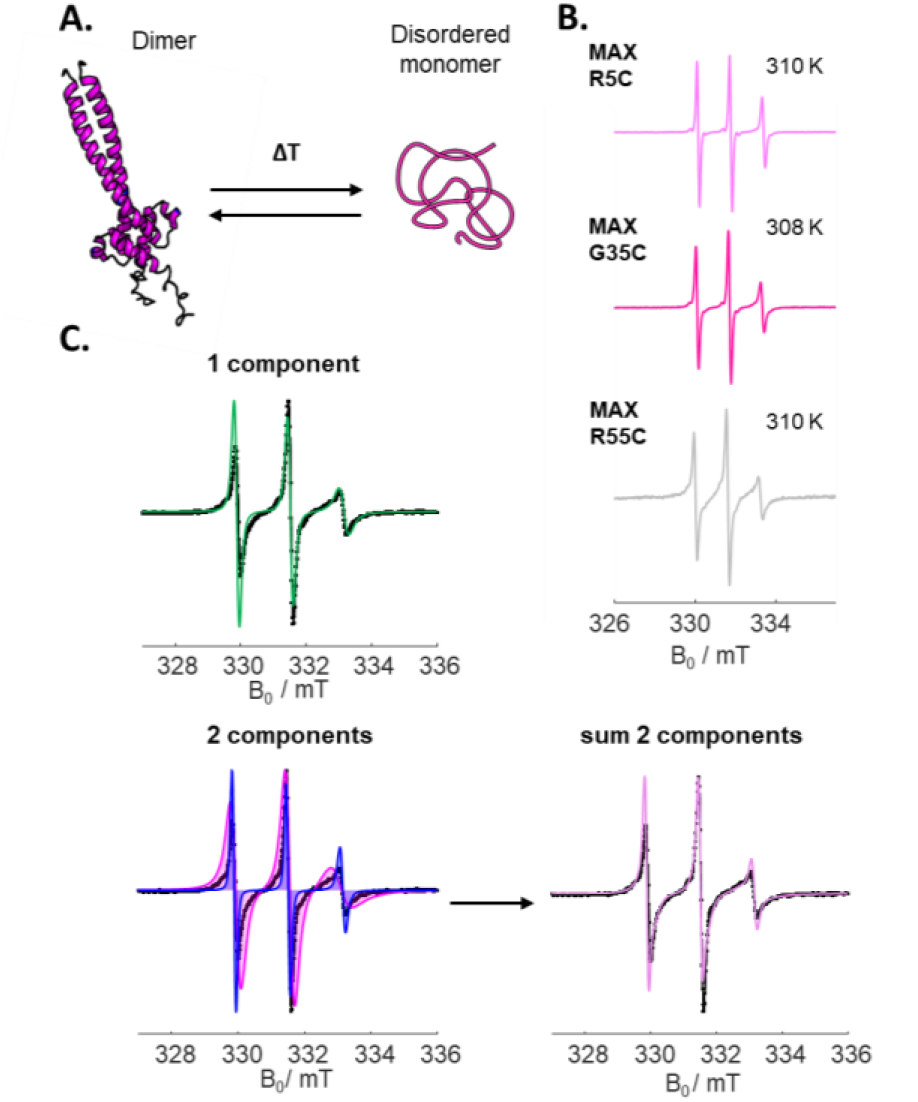
MAX behavior at high temperatures and EPR fitting profiles. A. Schematic representation of MAX dimer transition to the disordered monomer upon temperature increase. B. CW EPR profiles of MAX R5C, G35C, and R55C at temperatures higher than 300 K. The spectrum can be dissected into two components, one slow and one fast. C. One-component vs. two-component fit of the experimental data. The two-spin systems simulation (blue and magenta, which merges into the pink profile) better describes the experimental data in comparison to the single-component simulation (green profile).

To quantify the temperature-dependence, we fitted the weighted contributions of the two spectral components to sigmoidal functions (*Fig. 3*) and extracted their inflection points. These correspond to the temperatures at which the fast and slow components become equally populated, *i*.*e*., the cross-over point. The corresponding temperatures *T*_c_ are shown in the bar chart in Fig. 4 and listed Table 1.

**Table 1.** Tc (in *K*) derived from the EPR spectra simulations at different temperatures for the three mutants.

| Melting temperatures (K) |  |  |  |
| --- | --- | --- | --- |
| Mutation | MAX:MAX | MAX:MAX-DNA | MAX:MAX-DNA / BRCA1 |
| R5C | 305.1 | 322.5 | 313.6 |
| G35C | 312.1 | 323.1 | 321 |
| R55C | 321.7 | 321.3 | 323.5 |

**Figure 3.**
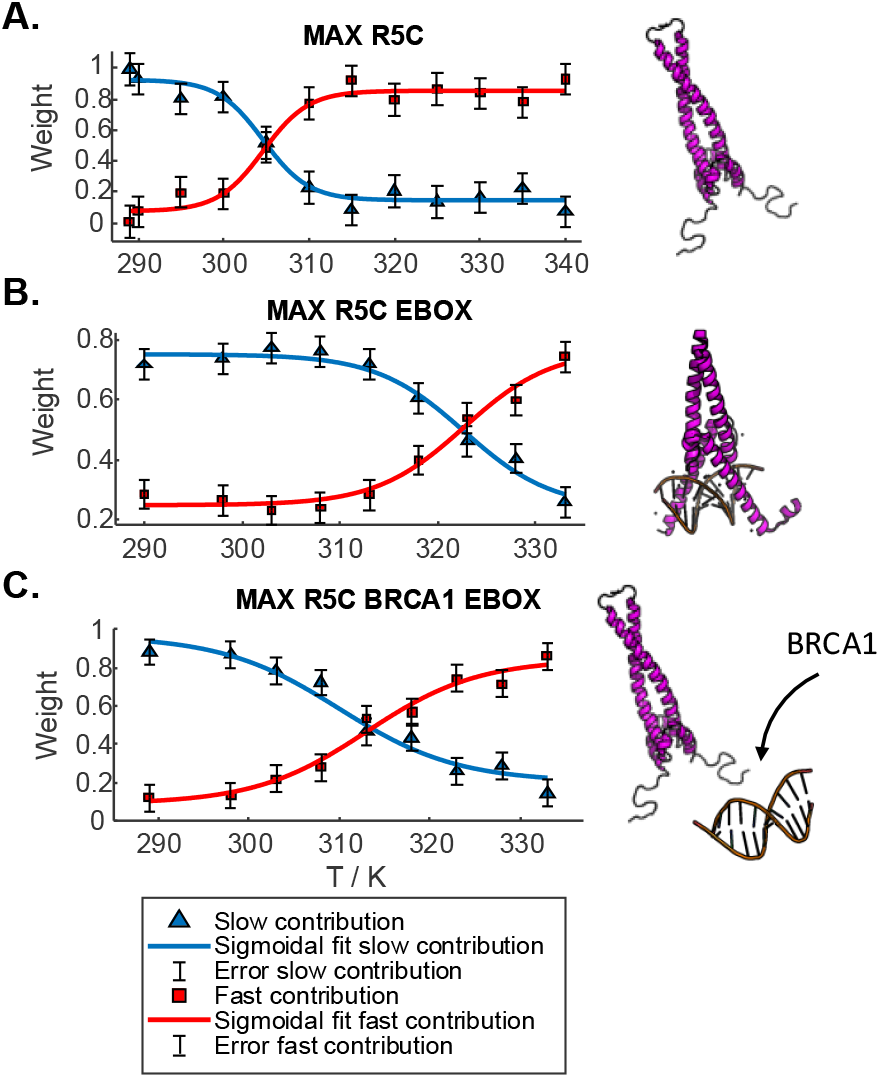
Sigmoidal fitting of fast (red squares) and slow (blue triangles) components’ weights at different temperatures for MAX R5C. The blue and red solid lines indicate the sigmoidal fitting. The panels represent the profiles of MAX R5C dimer (A.), MAX R5C dimer with EBOX (B.), and MAX R5C dimer with EBOX and BRCA1 (C.), respectively.

**Figure 4.**
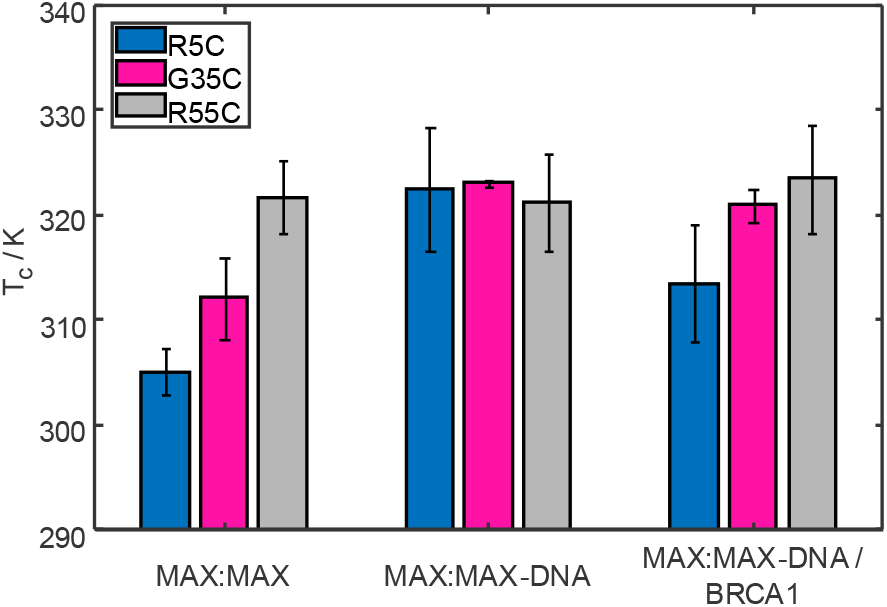
Comparison of melting temperatures of the three MAX with and without DNA and BRCA1. It is noticeable the important melting temperature difference between the NTD, HLH, and LZ regions of MAX:MAX; when the DNA is added, the dimer stability increases. In the presence of BRCA1, the original melting behavior of the dimer is partially restored.

We found that the NTD showed the lowest *T*_c_ around 305 K, followed by the HLH domain at 310 K and the LZ at 322 K. These temperatures correspond to the stability of the different MAX domains. The unfolded NTD with the most significant conformational plasticity transitions first. This is followed by the HLH, recently shown to be involved in large-scale dynamic fluctuations^15^, albeit with a stable secondary structure; lastly, the rigid LZ transitions. Thus, intuitively, these temperatures were attributed to the unfolding/dissociation of the individual MAX domains.

However, it is important to note that the underlying molecular processes—monomer-dimer exchange and local temperature-dependent MTSL dynamics etc.—are more complex, and the EPR-detected population shifts should be interpreted as ensemble-averaged reporters of these composite motions.

Nonetheless, the ratio between the slower and the faster components changes upon the dissociation of the MAX:MAX dimer and the concomitant formation of an intrinsically disordered monomeric state.^27^ Orthogonal NMR-based control experiments showed the same domain-specific destabilization patterns in the absence of spin labels, thereby ruling out label-induced artifacts or local probe dynamics as the primary cause of the observed temperature dependencies (Supporting Information Fig. S1).

### DNA Binding Shifts the Dissociation Profile of MAX Toward Cooperative Melting

We next compared these thermal profiles with data obtained in the presence of cognate DNA, the 5’-CACGTG-3’ enhancer box (EBOX) motif^6^, which has been known to bind the NTD of MAX efficiently.^15^ (*Fig. 1., C*. and *Fig. 2, B*.) Interestingly, for all three labeled positions, the equilibrium towards a faster motion was now found to be concerted at 322 K, corresponding to the cross-over point of the LZ label after DNA binding (*Fig. 3, B*.). Thus, a delayed transition for the NTD and HLH labels was observed, consistent with enhanced structural stabilization by DNA binding. Notably, only above 320 K did the fast component dominate the spectra, suggesting a sharp, cooperative transition (Supporting Information Fig. S2 to S4). This behavior contrasts with the gradual domain-by-domain unfolding seen in DNA-free conditions.

For R5C, *T*_c_ shifted from ~305 K (DNA-free) to ~323 K (DNA-bound) (*Fig. 4*). G35C showed a more modest stabilization (312 K *vs*. 323 K), while R55C exhibited near-identical melting behavior in both conditions (~322 K), indicating that LZ stability is largely independent of DNA. These results align with the structural model in which DNA binding rigidifies the NTD and HLH but not the distal LZ domain. The simultaneous dissociation of all labels in the MAX:MAX–DNA trimer, thus, appears to be initiated by destabilization of the LZ, which causes the entire complex to disassemble.

### BRCA1 Facilitates Dissociation by Destabilizing the DNA-Bound Complex

To test whether MAX:MAX–DNA dissociation can be facilitated by a third partner, we introduced a BRCA1 fragment (residues 219–504), previously shown to bind the EBOX DNA motif.^34^ Upon BRCA1^219–504^ addition, we observed pronounced shifts in the EPR temperature profiles and *T*_c_, particularly for the R5C and G35C mutants (*Fig. 3, C*., and *Fig. 4*). The presence of BRCA1 reduced the DNA-induced stabilization effect: *T*_c_ of R5C was elevated by ~8 K with EBOX DNA alone, but by only ~4 K when BRCA1 was also present. G35C exhibited a similar though less marked response. In contrast, R55C remained unaffected, suggesting that BRCA1 competes with MAX for DNA interaction rather than perturbing the LZ-mediated stability of the dimer.

These observations indicate that BRCA1 reduces the stabilizing influence of DNA on MAX, facilitating dissociation and allowing for earlier transitions into the dissociated states.

## Discussion

The *T*_c_ values provide an avenue for a mechanistic understanding of the temperature- and ligand-dependent dissociation behavior of the MAX:MAX dimer, particularly within its trimeric DNA-bound complex. In the absence of DNA, dissociation follows a domain-specific and sequential pathway: the intrinsically disordered N-terminal domain (NTD) exhibits early melting, followed by unfolding of the helix-loop-helix (HLH) domain, and finally the leucine zipper (LZ), which anchors the dimer. This stepwise disengagement reveals a modular structural hierarchy, consistent with a model in which local conformational fluctuations initiate dissociation prior to complete complex breakdown.

Strikingly, when MAX is bound to its cognate EBOX DNA, this modular dissociation is no longer observed. Instead, all thermally induced conformational changes converge into a single cooperative transition temperature. Our data demonstrate that this unified ‘melting’ event is governed by destabilization of the LZ. Once this key dimerization interface begins to unfold, the entire MAX:MAX-DNA trimer collapses. The synchronicity of this transition underscores a highly cooperative unbinding mechanism in the DNA-bound state, in which destabilization of the LZ acts as the critical trigger for dissociation of the whole trimeric complex.

The introduction of BRCA1^219–504^ destabilizes the MAX:MAX-DNA complex by competitively binding the DNA, effectively impairing the stabilizing influence of DNA on MAX dimer. As a result, domain-specific melting is present even in the presence of DNA, with early dissociation of the NTD and HLH domains preceding disruption of the LZ. This indicates that BRCA1 shifts the dissociation sequence such that DNA unbinding is facilitated before complete dissociation of the dimeric protein scaffold— evidence of facilitated dissociation.

The observed domain-resolved destabilization profile thereby provides direct evidence that BRCA1 facilitates transcription unbinding through a selective competition mechanism. Indeed, BRCA1 lowers the thermodynamic stability of DNA-bound MAX at the NTD and HLH regions, without perturbing the structural integrity of the LZ-mediated dimer. This implies that BRCA1 promotes MAX release from DNA by displacing it from the EBOX site, not by inducing dimer dissociation. Such domain-specific modulation is consistent with a facilitated dissociation mechanism and highlights the modularity of TF regulation.

## Conclusions

This study provides domain-resolved insights into the dissociation mechanism of the transcription factor MAX from its target DNA, revealing how the intrinsically disordered BRCA1^219–504^ fragment facilitates unbinding via competitive destabilization. Using site-directed spin labeling and continuous wave EPR spectroscopy, we captured discrete thermal transitions in the N-terminal and HLH domains of MAX, while the leucine zipper remained stable—demonstrating a clear sequence of domain-specific dissociation steps.

Our findings show that DNA binding enforces a highly cooperative melting behavior in the MAX:MAX–DNA trimeric complex, where destabilization of the leucine zipper triggers simultaneous collapse of all domains. The addition of BRCA1^219–504^, however, disrupts this coupling: by binding to the EBOX motif, it selectively reduces DNA-induced stabilization of the N-terminal and HLH regions, thereby promoting MAX dissociation without dismantling the dimer. This is a hallmark of facilitated dissociation, directly observed here for the first time with domain-level precision.

Crucially, this mechanistic dissection would be inaccessible with conventional competition assays, circular dichroism, or ensemble-based thermodynamic methods, which report only averaged unfolding or binding behavior. EPR spectroscopy offers a suite of advantages: it is intrinsically local, highly sensitive to changes in molecular motion, and can be performed in native solution environments at temperatures close to physiological. This allows detection of subtle nanoscale changes in protein dynamics that precede or accompany macroscopic dissociation events—providing a level of spatial and kinetic resolution not achievable with other techniques.

Our work establishes EPR spectroscopy as a powerful platform for probing facilitated dissociation mechanisms in transcriptional regulators. These findings offer a model for BRCA1-mediated regulation in the MYC:MAX axis and highlight a broadly applicable strategy for studying competitive unbinding in disordered or conformationally heterogeneous systems.

## Materials and methods

### Protein expression and sample preparation

The his-tagged MAX, and BRCA1 DNA codifying sequences were inserted in the vector pET21(a)+ and employed for Rosetta 2 *E. coli* cells transformation. The transformed bacteria were grown at 37° C up to an absorbance of 0.6 at 600 nm, and then resuspended in M9 (with homogeneously ^13^C labeled glucose 1 g/L and ^15^N ammonium chloride 1 g/L). The protein expression was then triggered by the addition of IPTG and conducted overnight at 30° C. For the full protein purification of BRCA1 and MAX, we followed the procedures reported in ^35^ and ^33^, respectively.

For the EPR experiments, we purchased three different pET21(a)+ vectors subcloned with the sequence codifying the three variants of MAX, R5C, G35C, and R55C. The expressed proteins were MTSL-tagged according to the reference ^36^. MTSL-tagged MAX mutants were buffer exchanged against MES buffer. Labelling efficiency was always above 98% as probed by DTNB assays.

We purchased the CACGTG DNA filaments forming the EBOX double-strand from Eurofins Austria.

### EPR

The Continuous Wave (CW) X-Band measurements have been recorded using a Bruker ElexSys E500 instrument (9.34 GHz) equipped with a super-high Q resonator (ER4122SHQE-W1). Samples (20 μL) with a protein concentration of 100 μM were loaded into a microcapillary tube. All measurements were recorded with 100-kHz field modulation. The temperature-stabilized measurements were performed using a temperature controller (ER 4131 VT, Bruker). The spectra were recorded with 0.25 G modulation amplitude for measurements in the range 290 K – 335 K.

All the raw CW EPR spectra are reported in the Supporting information, from SI5 to SI13.

The home-written MATLAB routine (script) for fitting of the room temperature CW EPR spectra can be found in the Supporting Information as Script S1.

### Simulations and Data Fitting

The EPR spectra were simulated with a homemade MATLAB 2019a script implementing the “chili” function of the EasySpin 5.7.0 package.

At <300 K the EPR spectra were successfully simulated by a single spin system for all the three mutants. At temperatures > 300 K, a good description of the experimental profiles was obtained by overlapping two spin systems with different *τ*_*c*_, representing the slow-and fast-motion contributions. The weights and *τ*_*c*_ obtained were varied during the fit of the spectra at different temperatures.

The weights of the slow and fast contributions over the temperature were then fitted in MATLAB with a sigmoidal function of equation: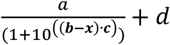.

### NMR

The H^N^-CON experiments were performed on a 600 MHz Avance Neo Bruker spectrometer equipped with a Bruker Prodigy TCI cryo-probe.

All samples were prepared in 25 mM MES buffer at pH 5.5. 10% D_2_O was added as lock solvent.

For the H^N^-CON selective ^1^H excitation and refocusing were achieved using PC9^37^ (bandwidth 6.25 ppm, duration 2001 µs), and REBURP^38^ (bandwidths 6.0-6.05 ppm, durations 1600-1614 µs) pulses, respectively. Selective proton pulses were centered at 8.3 ppm.

Selective ^13^C excitation and refocusing were achieved using Q5 (bandwidth 136 ppm, duration 300 µs), and Q3 (bandwidth 115 ppm, duration 199 µs) pulses.^39^ Selective ^13^C pulses were centered at 173 and 54 ppm for CO and Cα, respectively. IPAP acquisition was used to suppress CO-Cα couplings during acquisition. Non-selective ^15^N 90^°^ pulse durations were 32 µs. the number of scans was 32, the recycle delay was increased to 1 s, and the number of t_1_ increments collected was 256.

All the NMR analysis were performed using Topspin 4.0.7, CCPNMR and MATLAB R2019a. All data were apodized using 60° shifted sine bell functions before Fourier transform and baseline corrected using 5^th^-order polynomials.

## Supporting information

Supporting Information_Epasto et al.

## Acknowledgments

The authors acknowledge support from the NMR core facility of the Faculty of Chemistry, University Vienna. The project received funding from the European Research Council (ERC) under the European Union’s Horizon 2020 research and innovation programme (Grant agreement 801936). For the EPR experiments financial support from the IR INFRANALYTICS FR2054 is gratefully acknowledged. The authors thanks H. Ahouari (Chevreul Institute, Villeneuve d’Ascq) for technical assistance in recording CW spectra and troubleshooting.

## References

(1) Pang, Z. P.; Yang, N.; Vierbuchen, T.; Ostermeier, A.; Fuentes, D. R.; Yang, T. Q.; Citri, A.; Sebastiano, V.; Marro, S.; Südhof, T. C.; et al. Induction of human neuronal cells by defined transcription factors. Nature 2011, 476 (7359), 220–223. DOI: 10.1038/nature10202.

(2) Lambert, S. A.; Jolma, A.; Campitelli, L. F.; Das, P. K.; Yin, Y.; Albu, M.; Chen, X.; Taipale, J.; Hughes, T. R.; Weirauch, M. T. The Human Transcription Factors. Cell 2018, 172 (4), 650–665. DOI: 10.1016/j.cell.2018.01.029.

(3) Liu, J.; Perumal, N. B.; Oldfield, C. J.; Su, E. W.; Uversky, V. N.; Dunker, A. K. Intrinsic Disorder in Transcription Factors. Biochemistry 2006, 45 (22), 6873–6888. DOI: 10.1021/bi0602718.

(4) Bushweller, J. H. Targeting transcription factors in cancer — from undruggable to reality. Nature Reviews Cancer 2019, 19 (11), 611–624. DOI: 10.1038/s41568-019-0196-7.

(5) Wechsler, D. S.; Papoulas, O.; Dang, C. V.; Kingston, R. E. Differential Binding of C-Myc and Max to Nucleosomal DNA. Mol Cell Biol 1994, 14 (6), 4097–4107.

(6) Amati, B.; Land, H. Myc-Max-Mad: a transcription factor network controlling cell cycle progression, differentiation and death. Curr Opin Genet Dev 1994, 4 (1), 102–108.

(7) Brownlie, P.; Ceska, T. A.; Lamers, M.; Romier, C.; Stier, G.; Teo, H.; Suck, D. The crystal structure of an intact human Max-DNA complex: New insights into mechanisms of transcriptional control. Structure 1997, 5 (4), 509–520. DOI: Doi 10.1016/S0969-2126(97)00207-4.

(8) Nair, S. K.; Burley, S. K. X-Ray Structures of Myc-Max and Mad-Max Recognizing DNA: Molecular Bases of Regulation by Proto-Oncogenic Transcription Factors. Cell 2003, 112 (2), 193–205. DOI: 10.1016/S0092-8674(02)01284-9.

(9) Sauve, S.; Tremblay, L.; Lavigne, P. The NMR solution structure of a mutant of the max b/HLH/LZ free of DNA: Insights into the specific and reversible DNA binding mechanism of dimeric transcription factors. Journal of Molecular Biology 2004, 342 (3), 813–832. DOI: Doi 10.1016/J.Jmb.2004.07.058.

(10) Tu, W. B.; Helander, S.; Pilstal, R.; Hickman, K. A.; Lourenco, C.; Jurisica, I.; Raught, B.; Wallner, B.; Sunnerhagen, M.; Penn, L. Z. Myc and its interactors take shape. Bba-Gene Regul Mech 2015, 1849 (5), 469–483. DOI: 10.1016/j.bbagrm.2014.06.002.

(11) Amati, B.; Brooks, M. W.; Levy, N.; Littlewood, T. D.; Evan, G. I.; Land, H. Oncogenic Activity of the C-Myc Protein Requires Dimerization with Max. Cell 1993, 72 (2), 233–245. DOI: Doi 10.1016/0092-8674(93)90663-B.

(12) Macek, P.; Cliff, M. J.; Embrey, K. J.; Holdgate, G. A.; Nissink, J. W. M.; Panova, S.; Waltho, J. P.; Davies, R. A. Myc phosphorylation in its basic helix-loop-helix region destabilizes transient alpha-helical structures, disrupting Max and DNA binding. J Biol Chem 2018, 293 (24), 9301–9310. DOI: 10.1074/jbc.RA118.002709.

(13) Fieber, W.; Schneider, M. L.; Matt, T.; Kräutler, B.; Konrat, R.; Bister, K. Structure, function, and dynamics of the dimerization and DNA-binding domain of oncogenic transcription factor v-Myc11Edited by P. E. Wright. Journal of Molecular Biology 2001, 307 (5), 1395–1410. DOI: 10.1006/jmbi.2001.4537.

(14) Kamar, R. I.; Banigan, E. J.; Erbas, A.; Giuntoli, R. D.; Olvera de la Cruz, M.; Johnson, R. C.; Marko, J. F. Facilitated dissociation of transcription factors from single DNA binding sites. Proceedings of the National Academy of Sciences 2017, 114 (16), E3251–E3257.

(15) Sicoli, G.; Vezin, H.; Ledolter, K.; Kress, T.; Kurzbach, D. Conformational tuning of a DNA-bound transcription factor. Nucleic Acids Res 2019, 47 (10), 5429–5435. DOI: 10.1093/nar/gkz291.

(16) Tsai, M.-Y.; Zheng, W.; Chen, M.; Wolynes, P. G. Multiple binding configurations of Fis protein pairs on DNA: Facilitated dissociation versus cooperative dissociation. Journal of the American Chemical Society 2019, 141 (45), 18113–18126.

(17) Che, K.; Muttenthaler, M.; Kurzbach, D. Conformational selection of vasopressin upon V(1a) receptor binding. Comput Struct Biotechnol J 2021, 19, 5826–5833. DOI: 10.1016/j.csbj.2021.10.024.

(18) Igbaria-Jaber, Y.; Hofmann, L.; Gevorkyan-Airapetov, L.; Shenberger, Y.; Ruthstein, S. Revealing the DNA binding modes of CsoR by EPR spectroscopy. ACS omega 2023, 8 (42), 39886–39895.

(19) Dantu, S. C.; Khalil, M.; Bria, M.; Saint-Pierre, C.; Orio, M.; Gasparutto, D.; Sicoli, G. Cleaving DNA with DNA: Cooperative Tuning of Structure and Reactivity Driven by Copper Ions. Adv Sci (Weinh) 2024, 11 (16), e2306710. DOI: 10.1002/advs.202306710.

(20) Hofmann, L.; Mandato, A.; Saxena, S.; Ruthstein, S. The use of EPR spectroscopy to study transcription mechanisms. Biophysical Reviews 2022, 14 (5), 1141–1159.

(21) Yasin, A.; Mandato, A.; Hofmann, L.; Igbaria-Jaber, Y.; Shenberger, Y.; Gevorkyan-Airapetov, L.; Saxena, S.; Ruthstein, S. The Dynamic Plasticity of P. aeruginosa CueR Copper Transcription Factor upon Cofactor and DNA Binding. ChemBioChem 2024, 25 (15), e202400279.

(22) Casto, J.; Mandato, A.; Hofmann, L.; Yakobov, I.; Ghosh, S.; Ruthstein, S.; Saxena, S. Cu(ii)-based DNA labeling identifies the structural link between transcriptional activation and termination in a metalloregulator. Chemical Science 2022, 13 (6), 1693-1697, 10.1039/D1SC06563G. DOI: 10.1039/D1SC06563G.

(23) Roberts, M. G.; Dent, M. R.; Ramos, S.; Thielges, M. C.; Burstyn, J. N. Probing conformational dynamics of DNA binding by CO-sensing transcription factor, CooA. Journal of Inorganic Biochemistry 2024, 259, 112656. DOI: 10.1016/j.jinorgbio.2024.112656.

(24) Radman, K.; Crnolatac, I.; Bregović, N.; Matošević, Z. J.; Fernandes, P. A.; Merunka, D.; Žilić, D.; Piantanida, I.; Ašler, I. L.; Bertoša, B. Conformational change induced by binding of Mn2+ ions activates SloR transcription factor in Streptococcus mutans. International Journal of Biological Macromolecules 2025, 290, 138828. DOI: 10.1016/j.ijbiomac.2024.138828.

(25) Tang, S.; Hicks, N. D.; Cheng, Y.-S.; Silva, A.; Fortune, S. M.; Sacchettini, J. C. Structural and functional insight into the Mycobacterium tuberculosis protein PrpR reveals a novel type of transcription factor. Nucleic Acids Research 2019, 47 (18), 9934–9949. DOI: 10.1093/nar/gkz724 (acccessed 6/5/2025).

(26) Vancraenenbroeck, R.; Hofmann, H. Occupancies in the DNA-Binding Pathways of Intrinsically Disordered Helix-Loop-Helix Leucine-Zipper Proteins. J Phys Chem B 2018, 122 (49), 11460–11467. DOI: 10.1021/acs.jpcb.8b07351.

(27) Kizilsavas, G.; Ledolter, K.; Kurzbach, D. Hydrophobic Collapse of the Intrinsically Disordered Transcription Factor Myc Associated Factor X. Biochemistry 2017, 56 (40), 5365–5372. DOI: 10.1021/acs.biochem.7b00679.

(28) Sicoli, G.; Kress, T.; Vezin, H.; Ledolter, K.; Kurzbach, D. A Switch between Two Intrinsically Disordered Conformational Ensembles Modulates the Active Site of a Basic-Helix-Loop-Helix Transcription Factor. J Phys Chem Lett 2020, 11 (21), 8944–8951. DOI: 10.1021/acs.jpclett.0c02242.

(29) Schweiger, A.; Jeschke, G. Principles of pulse electron paramagnetic resonance; Oxford university press, 2001.

(30) Weil, J. A.; Bolton, J. R. Electron paramagnetic resonance: elementary theory and practical applications; John Wiley & Sons, 2007.

(31) Lavigne, P.; Crump, M. P.; Gagné, S. M.; Hodges, R. S.; Kay, C. M.; Sykes, B. D. Insights into the mechanism of heterodimerization from the 1H-NMR solution structure of the c-Myc-Max heterodimeric leucine zipper11Edited by P. E. Wright. Journal of Molecular Biology 1998, 281 (1), 165–181. DOI: 10.1006/jmbi.1998.1914.

(32) Sauve, S.; Naud, J. F.; Lavigne, P. The mechanism of discrimination between cognate and non-specific DNA by dimeric b/HLH/LZ transcription factors. J Mol Biol 2007, 365 (4), 1163–1175. DOI: 10.1016/j.jmb.2006.10.044.

(33) Epasto, L. M.; Che, K.; Kozak, F.; Selimovic, A.; Kaderavek, P.; Kurzbach, D. Toward protein NMR at physiological concentrations by hyperpolarized water-Finding and mapping uncharted conformational spaces. Sci Adv 2022, 8 (31), eabq5179. DOI: 10.1126/sciadv.abq5179.

(34) Che, K.; Kress, T.; Górka, M.; Žerko, S.; Kozminski, W.; Kurzbach, D. Coupled MD simulations and NMR reveal that the intrinsically disordered domain of the breast-cancer susceptibility 1 protein (BRCA1) binds head-on to DNA double-strand ends. Journal of Magnetic Resonance Open 2022, 12, 100069.

(35) Che, K.; Kress, T.; Górka, M.; Žerko, S.; Kozminski, W.; Kurzbach, D. Coupled MD simulations and NMR reveal that the intrinsically disordered domain of the breast-cancer susceptibility 1 protein (BRCA1) binds head-on to DNA double-strand ends. Journal of Magnetic Resonance Open 2022, 12-13, 100069. DOI: 10.1016/j.jmro.2022.100069.

(36) Kurzbach, D.; Platzer, G.; Schwarz, T. C.; Henen, M. A.; Konrat, R.; Hinderberger, D. Cooperative unfolding of compact conformations of the intrinsically disordered protein osteopontin. Biochemistry 2013, 52 (31), 5167–5175.

(37) Kupce, E.; Freeman, R. Wideband Excitation with Polychromatic Pulses. J. Magn. Reson. A 1994, 108, 268–273.

(38) Geen, H.; Freeman, R. Band-selective radiofrequency pulses. J. Magn. Reson. 1991, 93, 93–141.

(39) Emsley, L.; Bodenhausen, G. Gaussian pulse cascades: New analytical functions for rectangular selective inversion and in-phase excitation in NMR. Chem. Phys. Lett. 1990, 165 (469-476).

