## Supporting Information_Epasto et al. for "Domain-Specific EPR Spectroscopy to Monitor Facilitated Dissociation of a DNA-Transcription Factor Complex"

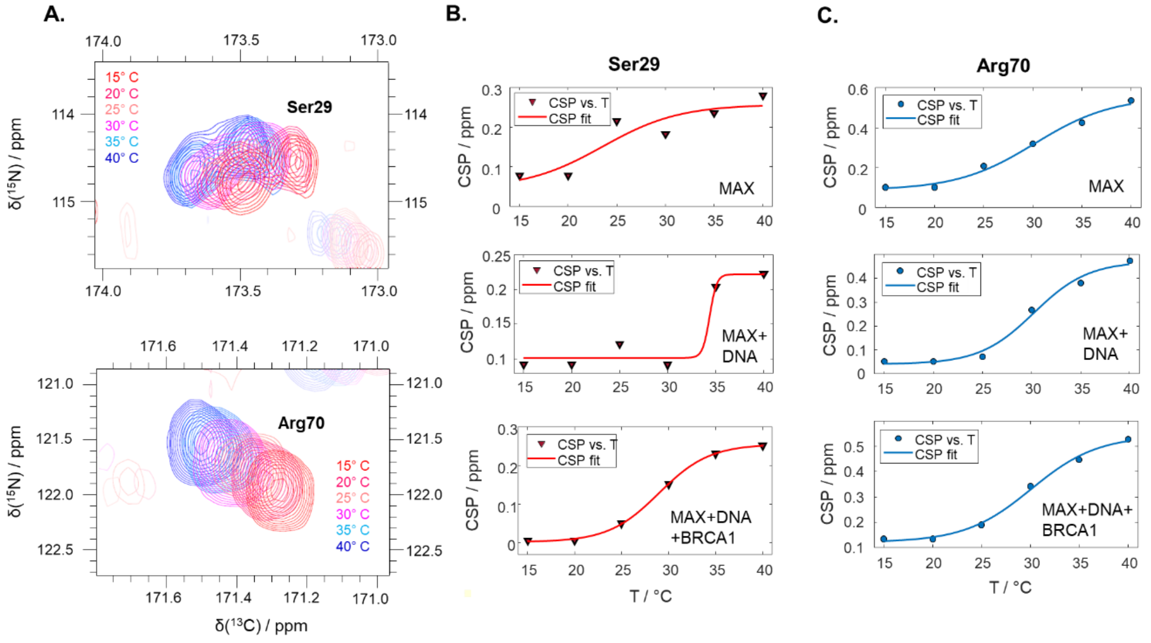


**Figure SI 1.** NMR $^{13}C$-$^{15}N$ correlation control experiments. A. NMR spectra overlap collected at temperatures between 15° C (in red) and 40° C (in blue) of two model amino acids, Serine 29 and Arginine 70, located in the helix 1 and in the leucine zipper region, respectively. B. and C. Chemical shift perturbation (CSP) profiles of Ser29 (B.) and Arg70 (C.) at different temperatures, modeled with a sinusoidal function (solid line). The melting point of MAX:MAX is identified as the inflection point of the sinusoidal fit. In the presence of DNA, Ser29 is stabilized; when BRCA1 is added to MAX:MAX-DNA, the CSP profile resembles the one of the Ser29 of MAX:MAX without DNA. Instead, Arg70 CSP profiles describe a slight stabilization upon the addition of DNA, but the peak shift is not heavily influenced by the addition of BRCA1.

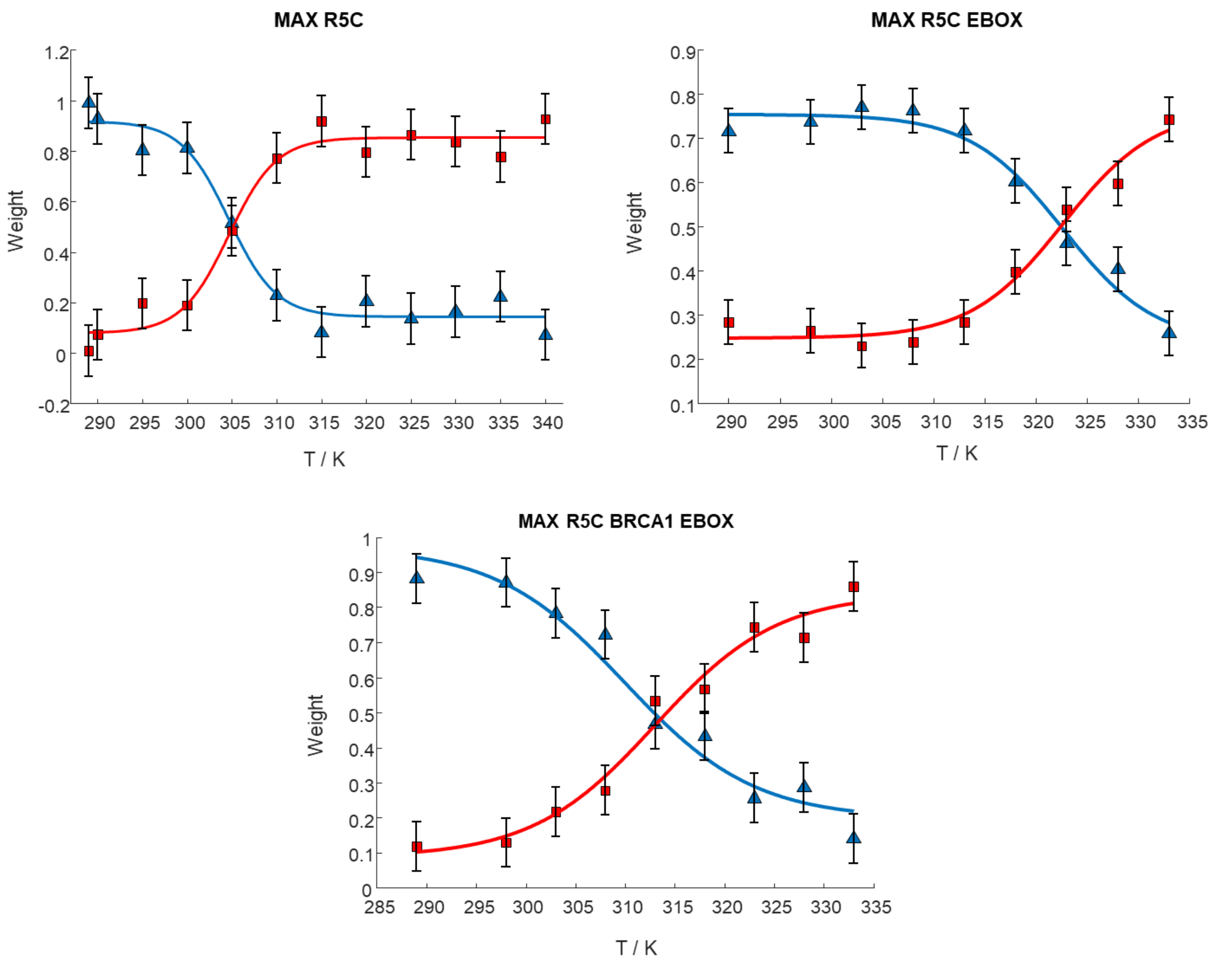


**Figure SI 2.** Weights of the slow contribution (blue triangles) and fast contribution (red squares) simulated from the CW EPR spectra of MAX R5C collected at different temperatures. The fit of the experimental data was performed with a sigmoidal function (blue and red lines, respectively). The melting temperature is identified as the crossing point between the fits. The experiments were performed in absence and presence of BRCA1 and/or EBOX DNA.

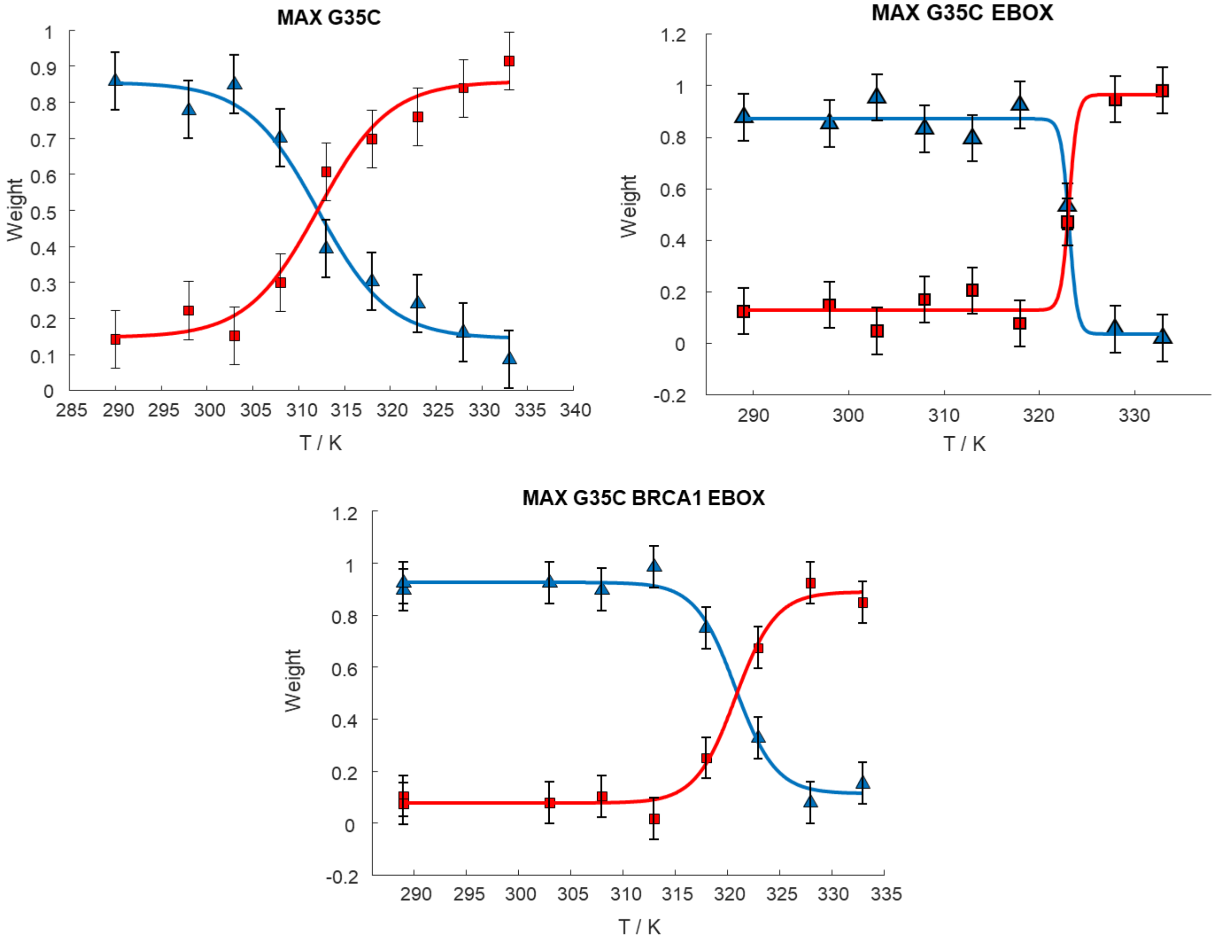


**Figure SI 3.** Weights of the slow contribution (blue triangles) and fast contribution (red squares) simulated from the CW EPR spectra of MAX G35C collected at different temperatures. The fit of the experimental data was performed with a sigmoidal function (blue and red lines, respectively). The melting temperature is identified as the crossing point between the fits. The experiments were performed in absence and presence of BRCA1 and/or EBOX DNA.

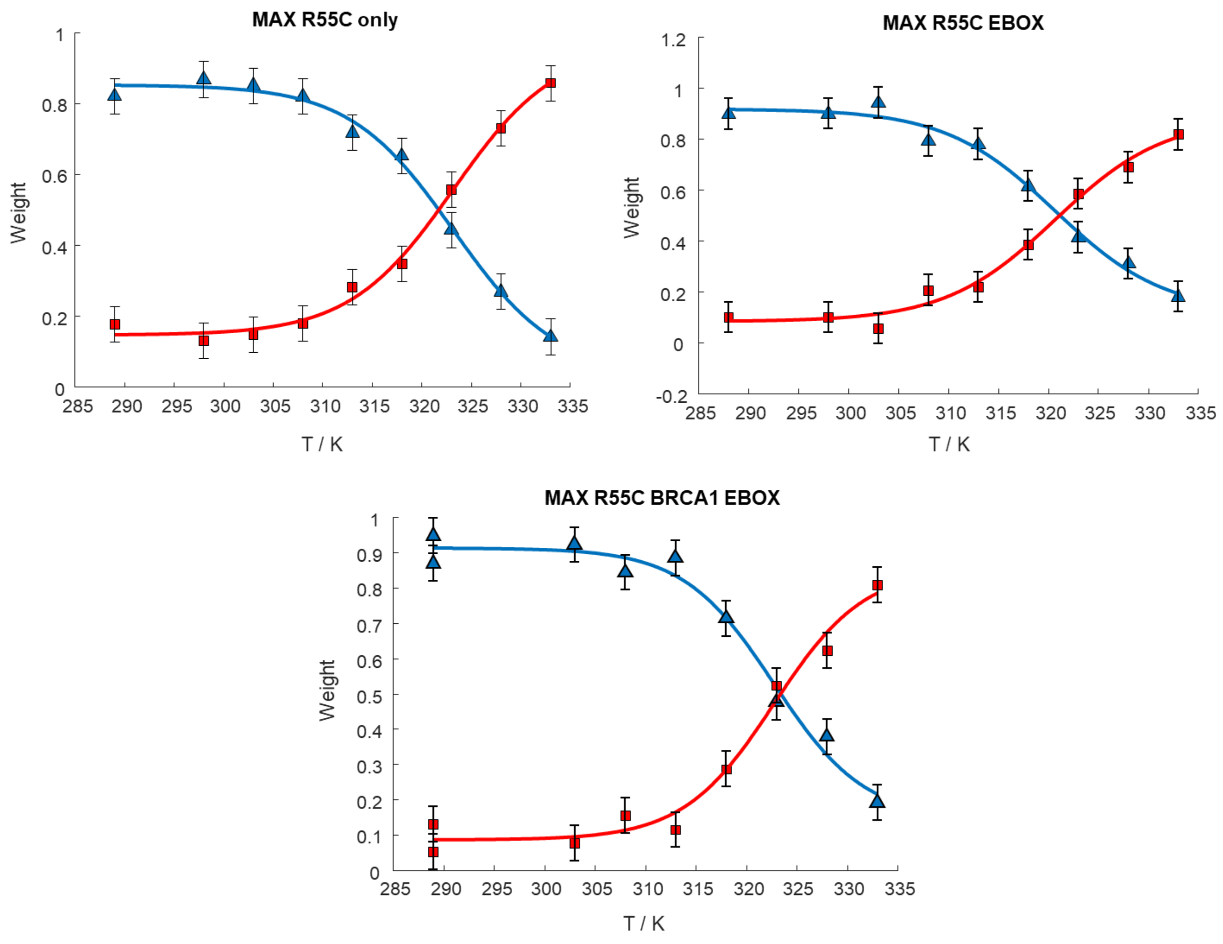


**Figure SI 4.** Weights of the slow contribution (blue triangles) and fast contribution (red squares) simulated from the CW EPR spectra of MAX R55C collected at different temperatures. The fit of the experimental data was performed with a sigmoidal function (blue and red lines, respectively). The melting temperature is identified as the crossing point between the fits. The experiments were performed in absence and presence of BRCA1 and/or EBOX DNA.

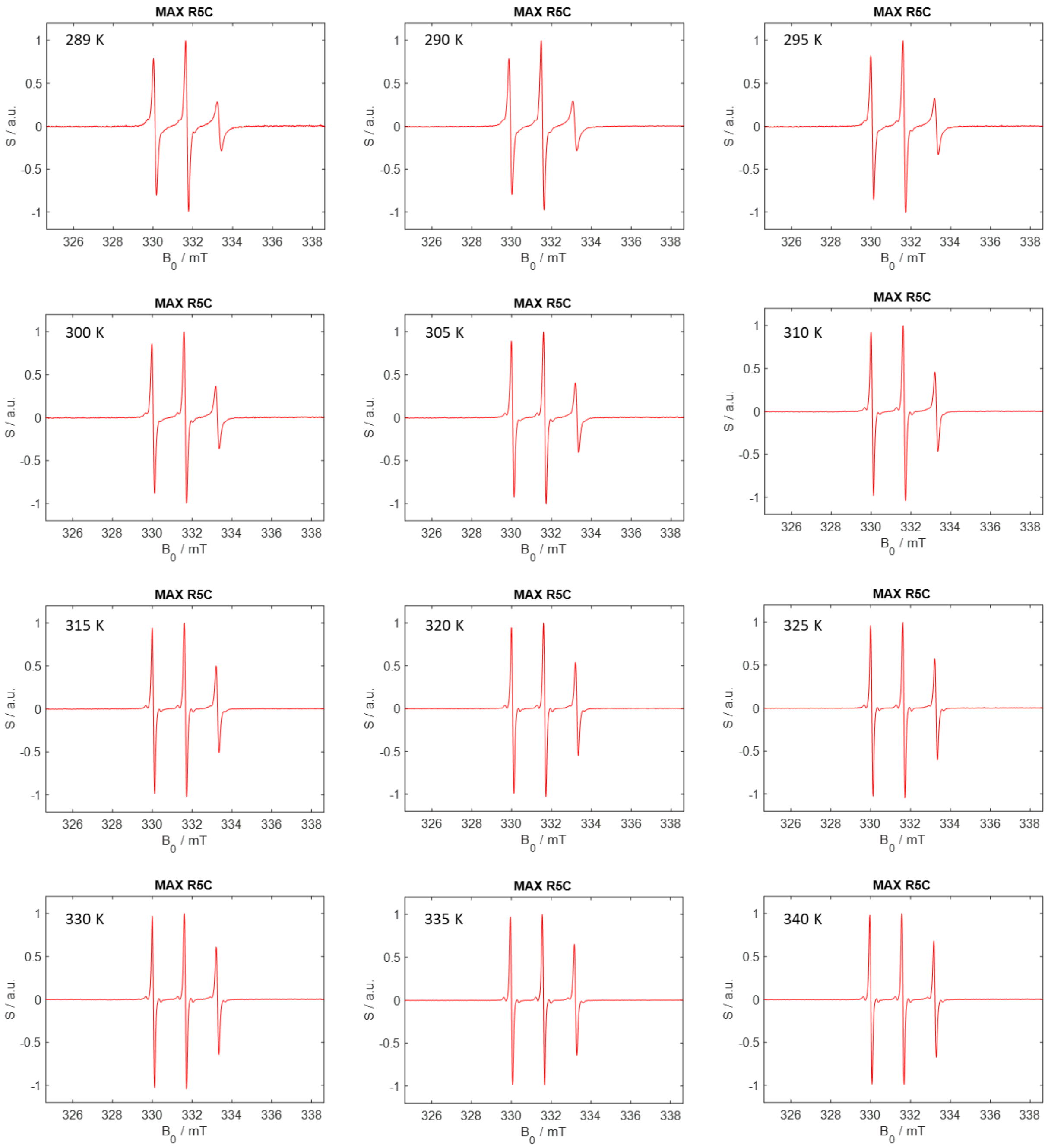


**Figure SI 5.** CW EPR spectra of MAX R5C. The experimental temperatures are indicated in the pictures.

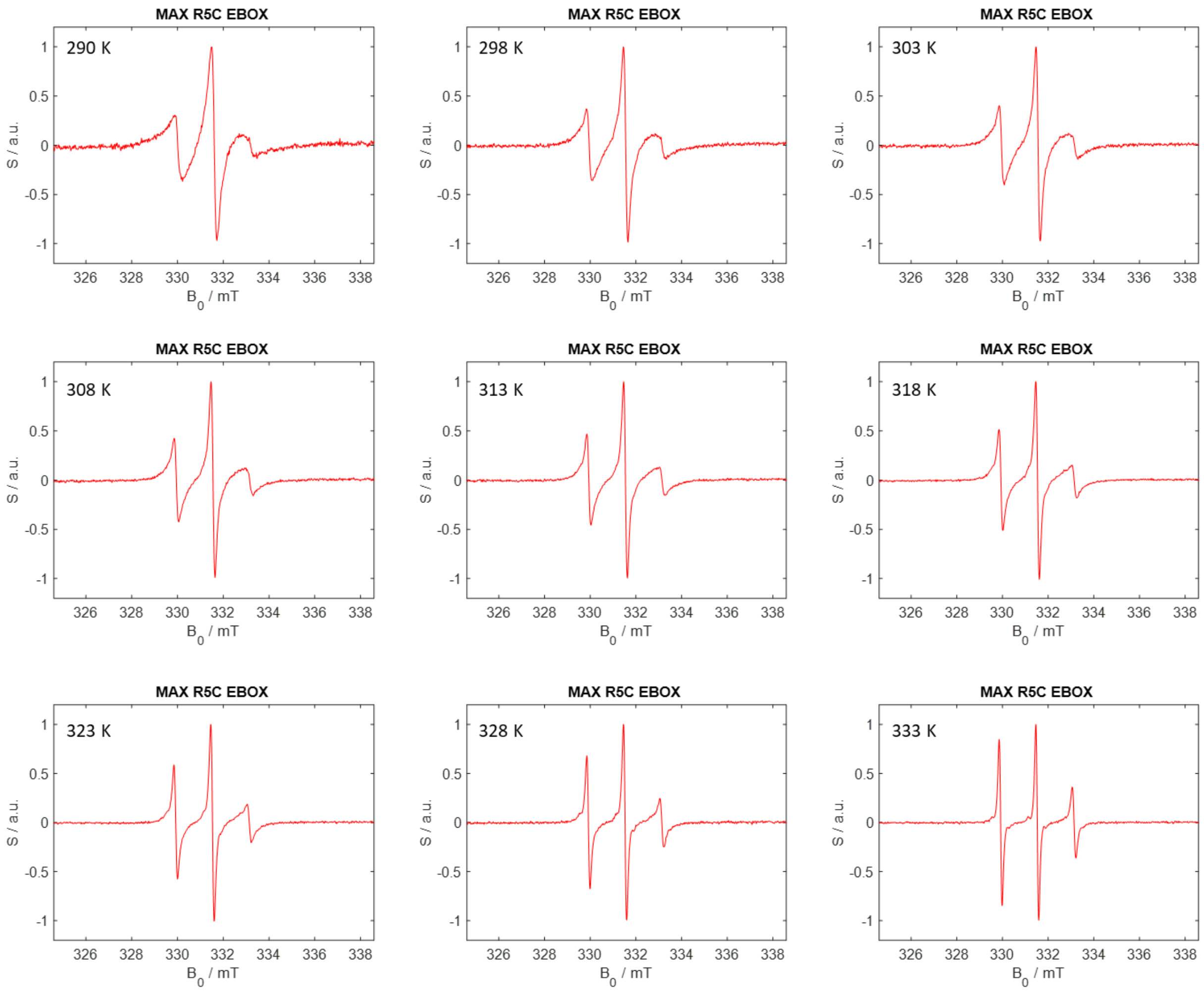


**Figure SI 6.** CW EPR spectra of MAX R5C + EBOX DNA. The experimental temperatures are indicated in the pictures.

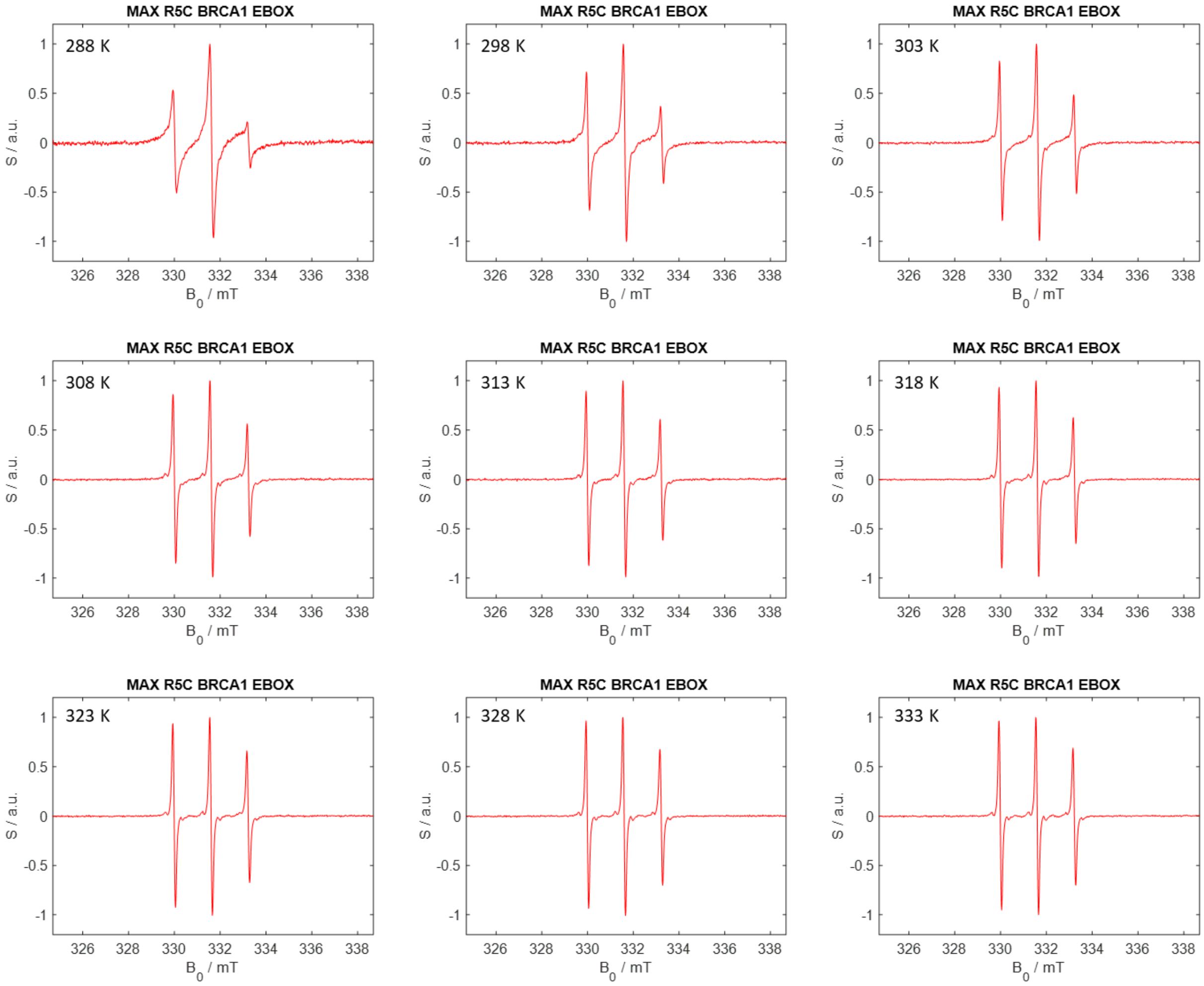


**Figure SI 7.** CW EPR spectra of MAX R5C + EBOX DNA + BRCA1. The experimental temperatures are indicated in the pictures.

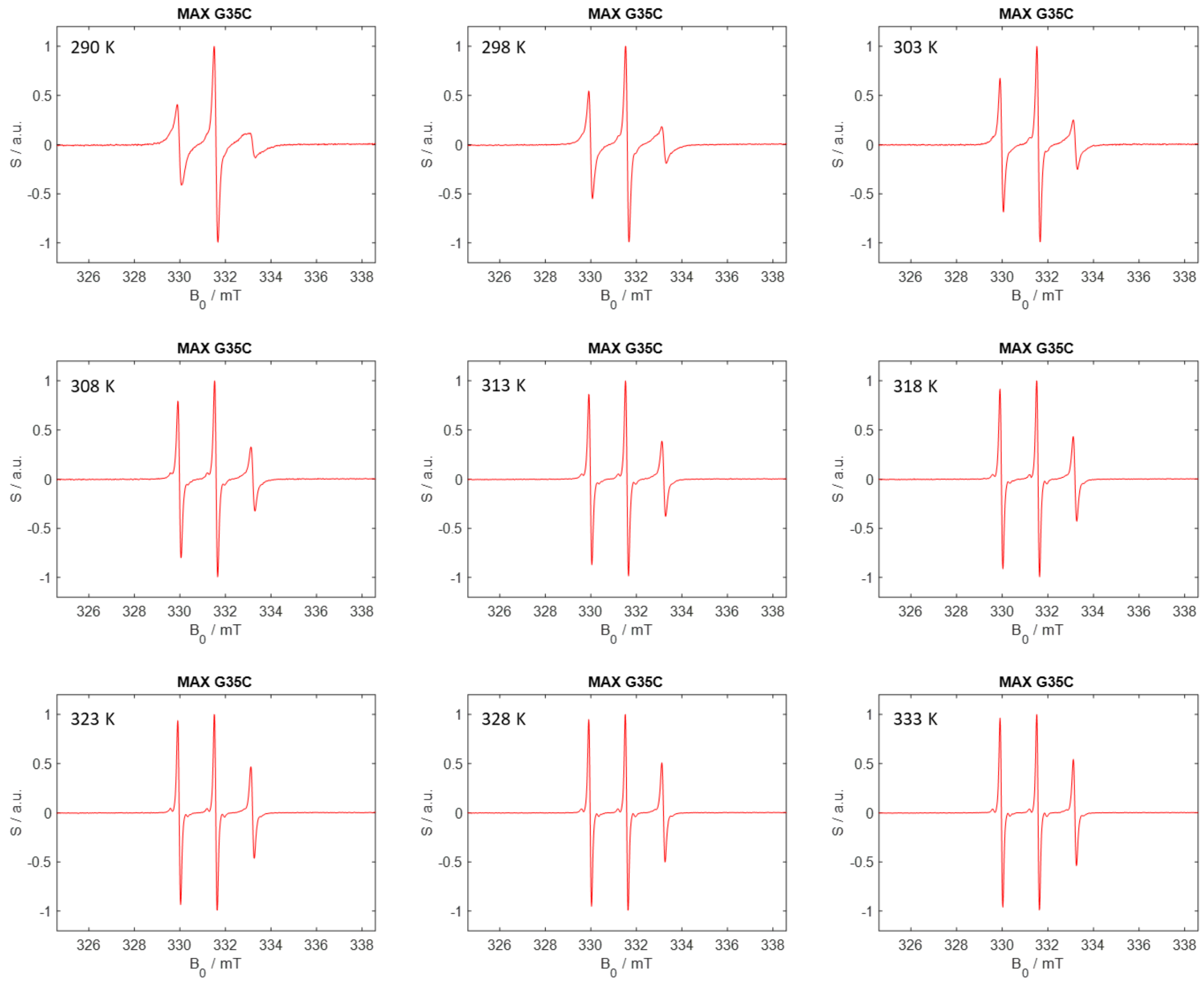


**Figure SI 8.** CW EPR spectra of MAX G35C. The experimental temperatures are indicated in the pictures.

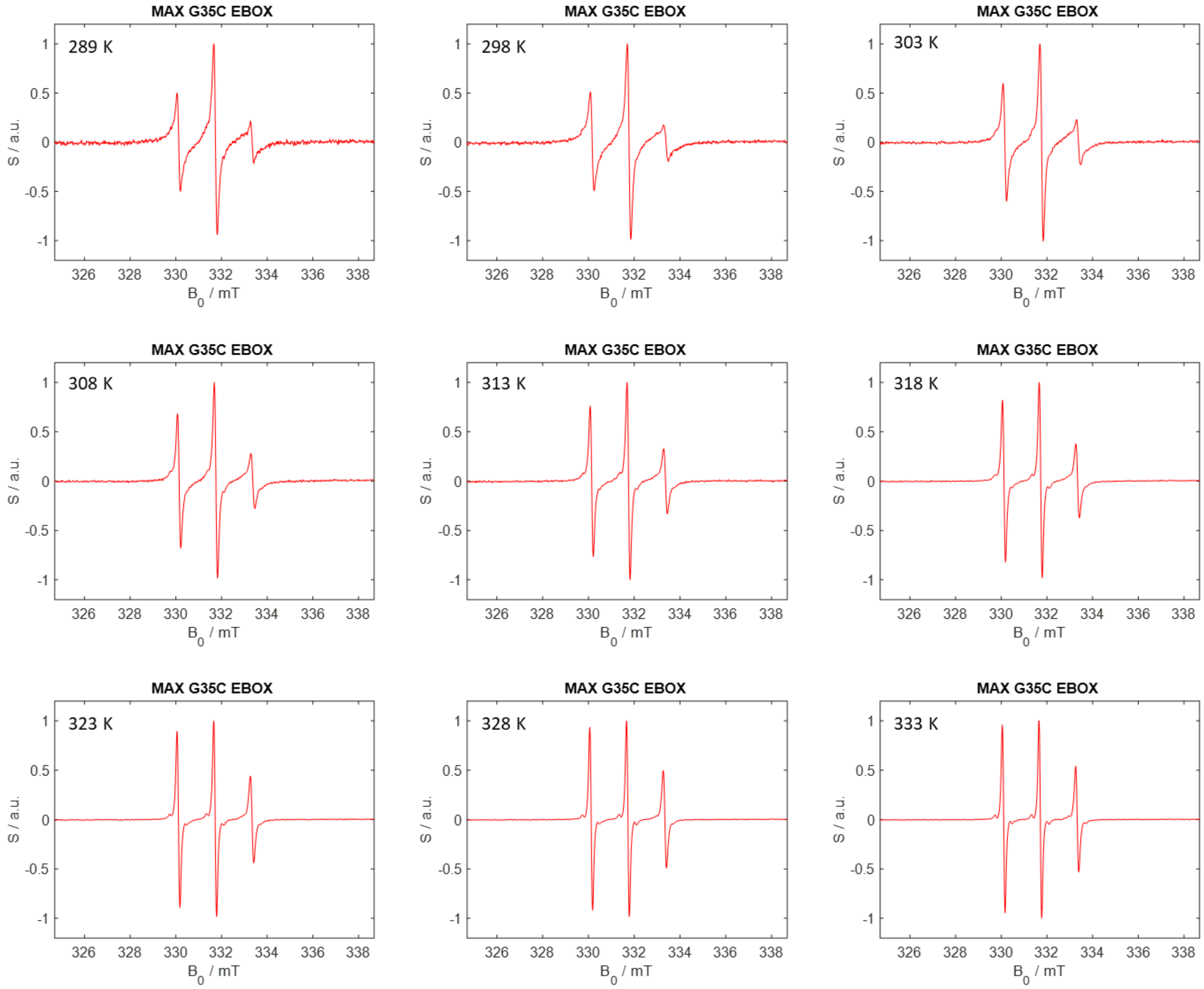


**Figure SI 9.** CW EPR spectra of MAX G35C + EBOX DNA. The experimental temperatures are indicated in the pictures.

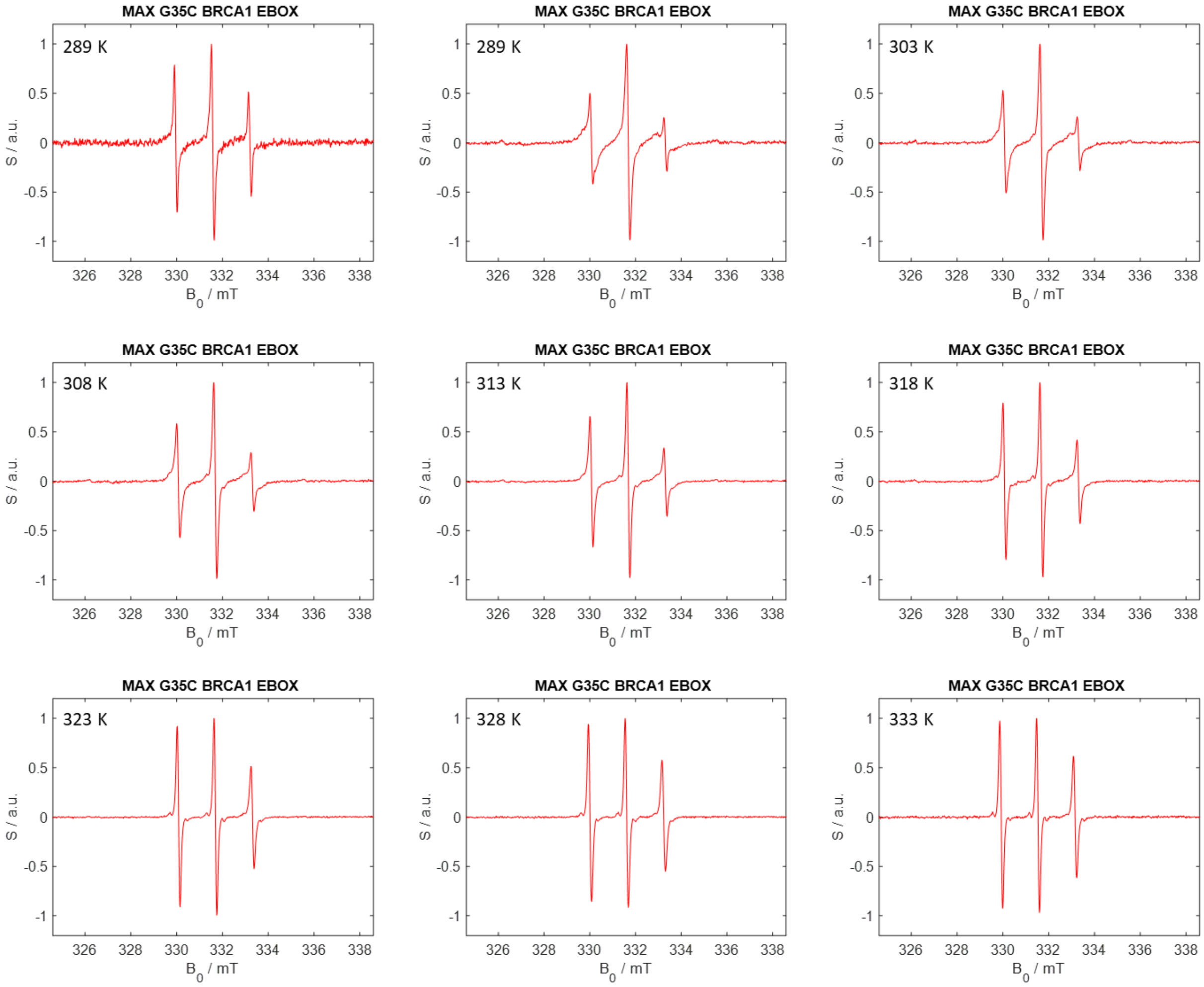


**Figure SI 10.** CW EPR spectra of MAX G35C + EBOX DNA + BRCA1. The experimental temperatures are indicated in the pictures.

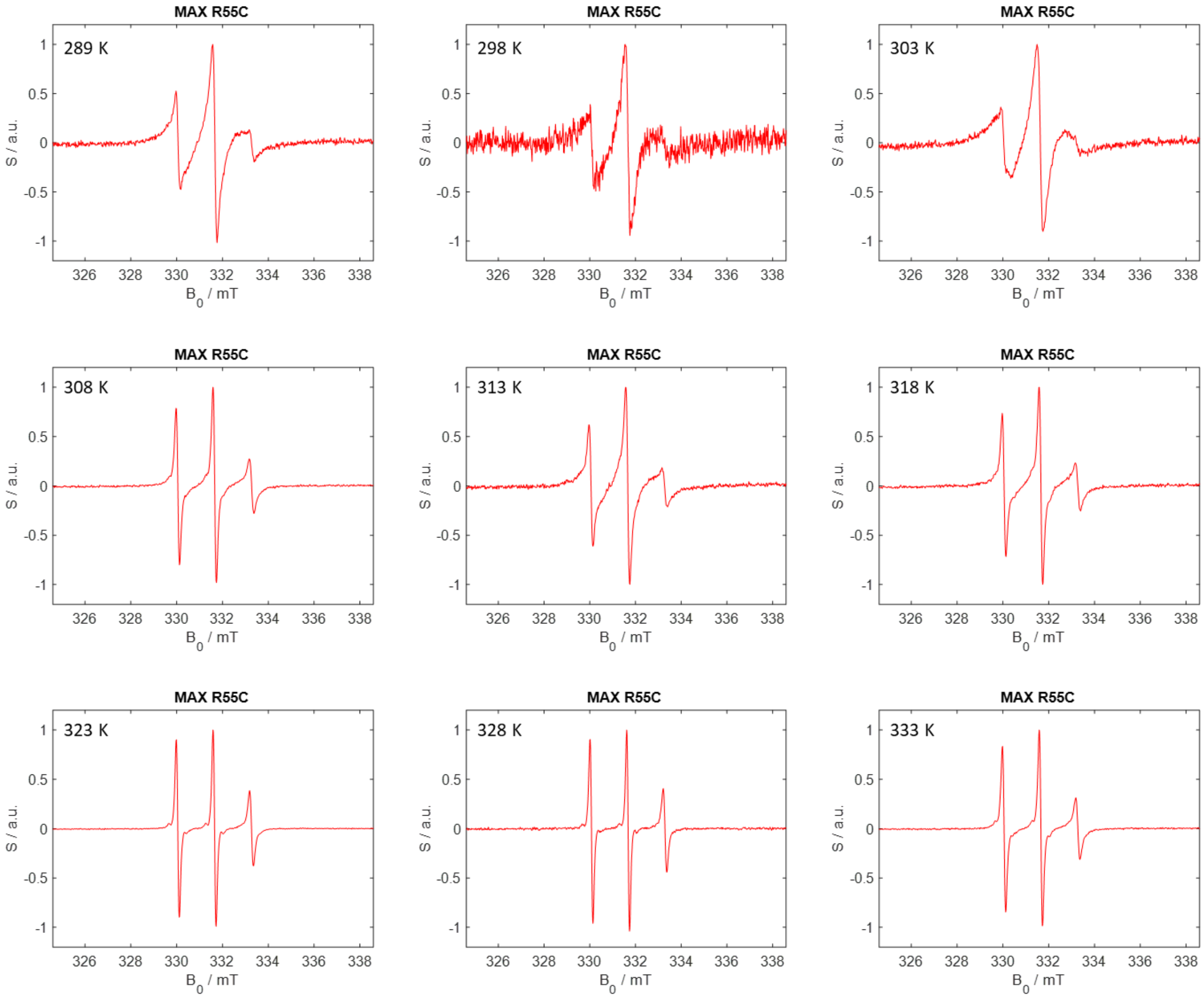


**Figure SI 11.** CW EPR spectra of MAX R55C. The experimental temperatures are indicated in the pictures.

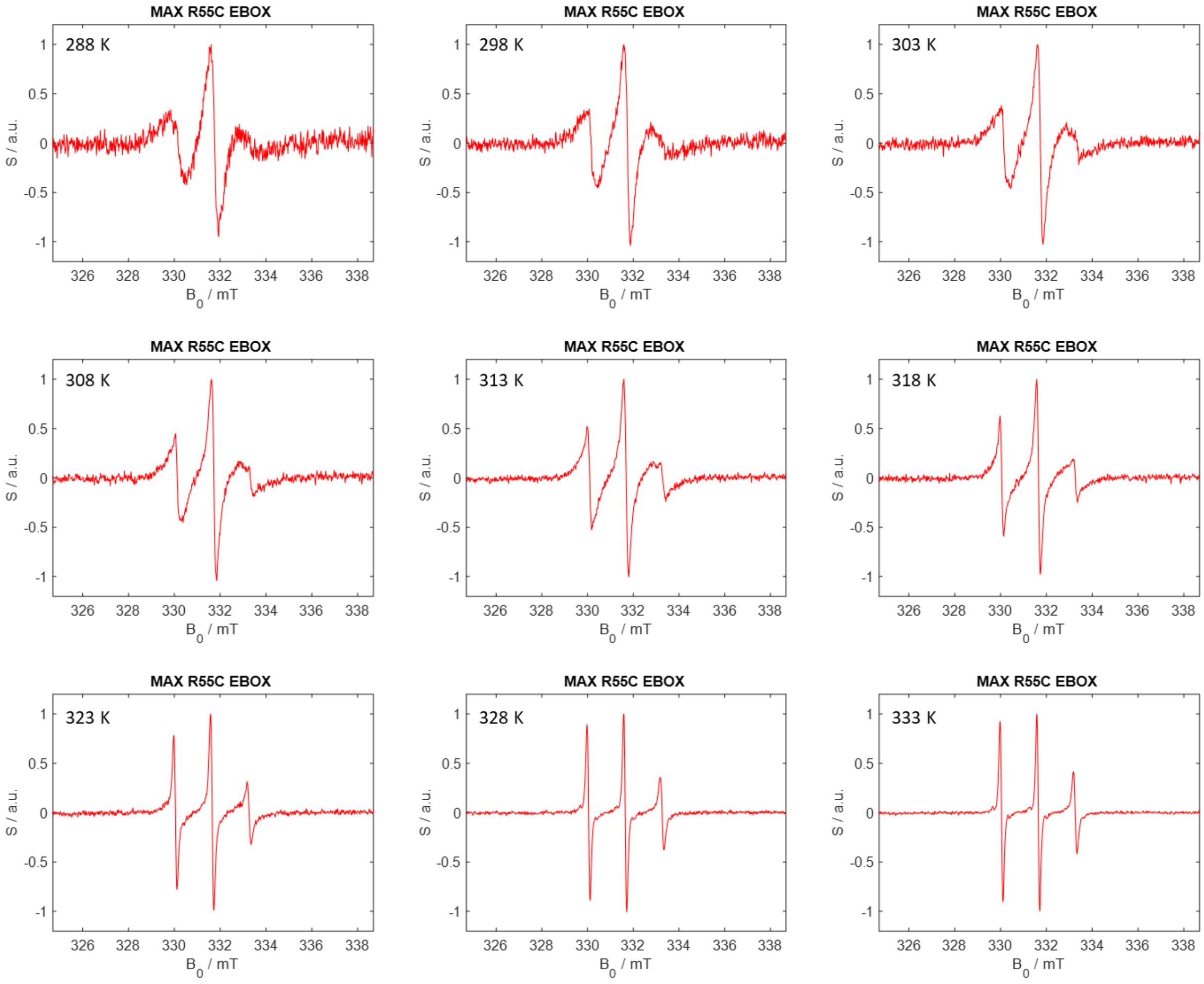


**Figure SI 12.** CW EPR spectra of MAX R55C + EBOX DNA. The experimental temperatures are indicated in the pictures.

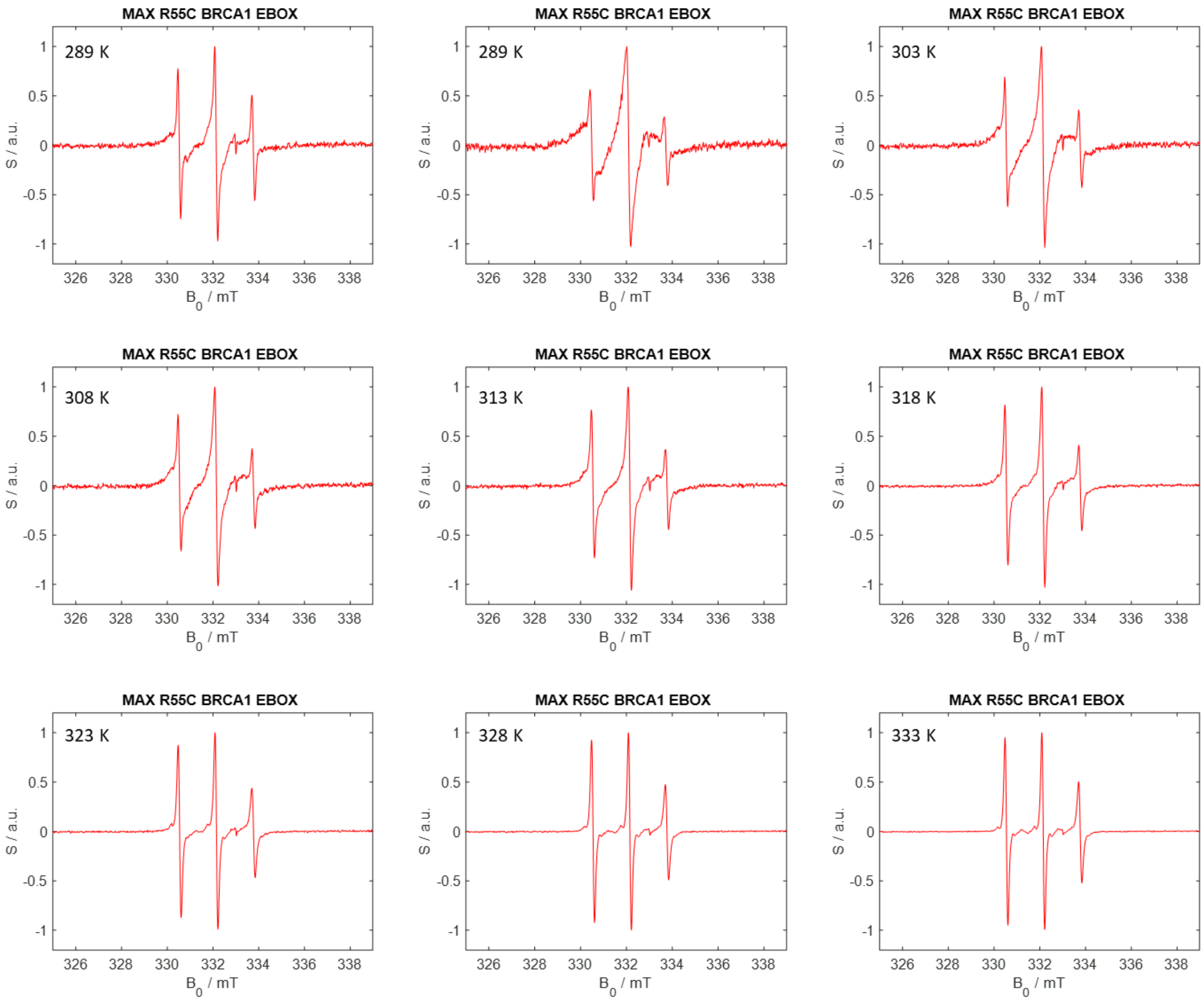


**Figure SI 13.** CW EPR spectra of MAX R55C + EBOX DNA + BRCA1. The experimental temperatures are indicated in the pictures.

**Matlab Script S1**

```
% Matlab script to simulate the EPR spectrum of TEMPOL
% using the EasySpin simulation package (www.easyspin.org)


%% simulate data

npts = 1024;

%% Spectrum

Exp.Harmonic = 1;
Exp.Temperature = 290;
Exp.mwFreq = str2double(par.FrequencyMon(1:5));
Exp.CenterSweep = [str2double(par.CenterField(1:4))/10
str2double(par.SweepWidth(1:3))/10];
Exp.Range = [B0(1,1)*10^-1 B0(end,1)*10^-1];
tcor1 = 1*10^-9;
tcor2 = 3*10^-10;

Sys0.Nucs = '14N';
Sys0.S = 1/2; %green
Sys0.lwpp=[0.1 0];
Sys0.g = [g1 g2 g3];
Sys0.A = [A1 A2 A3];
Sys0.logtcorr = log10(tcor1); %axial
Sys0.weight=0.9;

Sys1.Nucs = '14N';
Sys1.S = 1/2; % magenta
Sys1.lwpp=[0.08 0];
Sys1.g = [g1 g2 g3];
Sys1.A = [A1 A2 A3];
Sys1.logtcorr = log10(tcor2); %axial
Sys1.weight=0.1;

% Vary1.g = [0.01 0.01 0.01];
% Vary1.A = [1 10 1];
Vary1.logtcorr = 0.05;
% Vary1.lwpp=[0.05 0];
% Vary2.g = [0.01 0.01 0.01];
% Vary2.A = [1 10 1];
Vary2.logtcorr = 0.5;
% Vary2.lwpp=[0.1 0];

Vary1.weight=0.2;
Vary2.weight=0.2;

B0 = B0(:,5);
spec = data_norm(:,2);
Sys = {Sys0 };
Vary = {Vary1 };
```

```
esfit(spec,@chili,{Sys,Exp},{Vary});


plot(B0,fit1.fit);
eprsave('fitSpec',B0,fit1.fit,'EasySpin simulation');
%
data = [B0 fit1.fit];
save('fitSpec_ascii.txt','data','-ascii');
```
